## Supporting information for "A unified mechanism for innate and learned visual landmark guidance in the insect central complex"

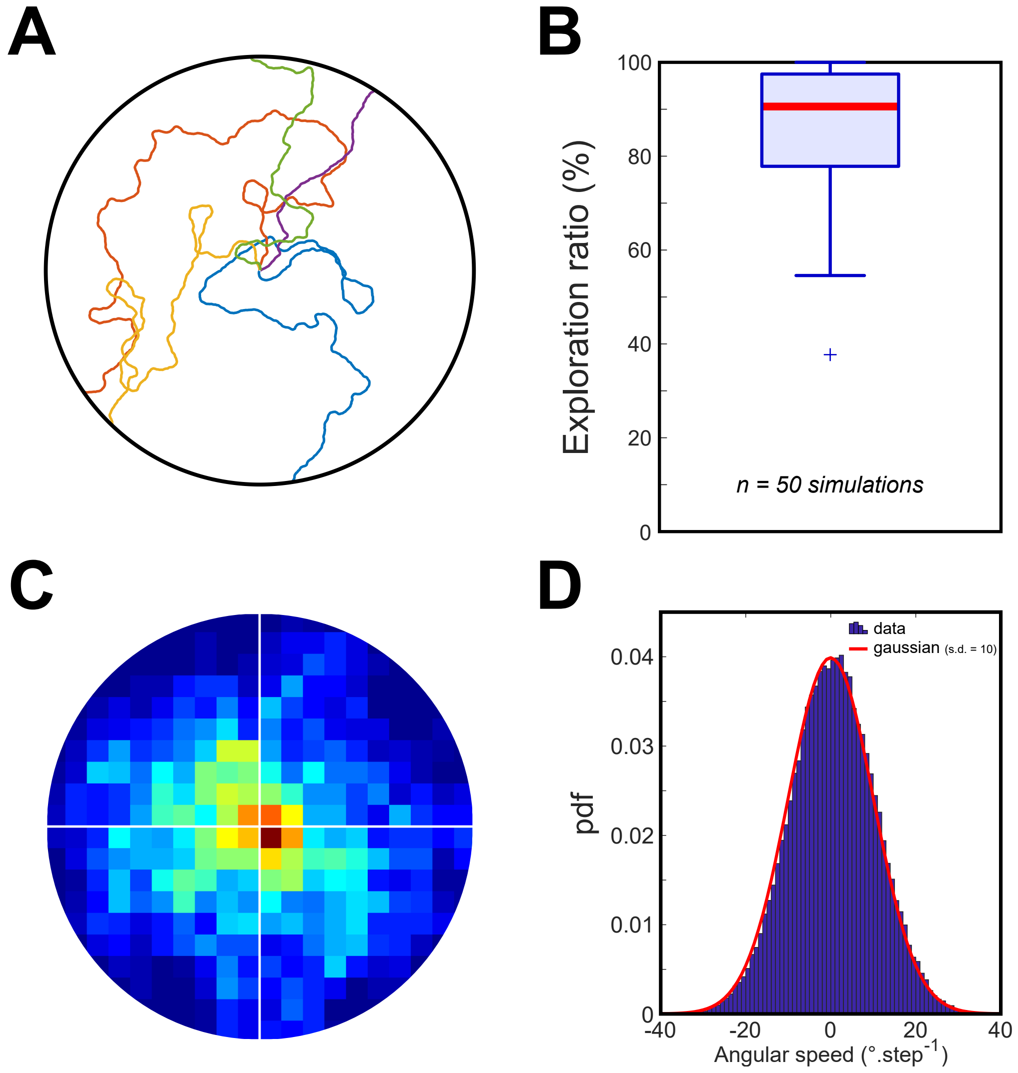


Figure S.1: **Simulations with no modulation of the EPG-PFL synapses.**
**A.** 5 examples of paths where the EPG-PFL3 synapse weights are set equal and constant.
**B.** Boxplot (Median: Red, Inter-quartile: shaded box) of the exploration ratio, calculated as the percentage of the 360° surroundings faced by the agent, during 50 simulations.
**C.** Heatmap of 50 simulations stacked adjusted to the cue location (0°).
**D.** Probability density function of the angular speed (all 50 simulations data stacked) corresponding to the 10° s.d. gaussian noise applied to the output of the steering model (red line).

| PreN | PreN ID | PostN | PostN ID | #syn |
|---|---|---|---|---|
| EPG R4 | 633546217 | PFL3 R3 | 1008028537 | 24 |
| EPG R3 | 478375456 | PFL3 R3 | 1008028537 | 15 |
| EPG R3 | 541118908 | PFL3 R3 | 1008028537 | 15 |
| EPG R4 | 695956656 | PFL3 R3 | 1008028537 | 8 |
| EPG R4 | 416642425 | PFL3 R3 | 1008028537 | 18 |
| EPG R3 | 5813080838 | PFL3 R3 | 1008028537 | 12 |
| EPG R4 | 416642425 | PFL3 R4 | 942172835 | 23 |
| EPG R3 | 478375456 | PFL3 R4 | 942172835 | 2 |
| EPG R3 | 541118908 | PFL3 R4 | 942172835 | 2 |
| EPG R4 | 695956656 | PFL3 R4 | 942172835 | 17 |
| EPG R4 | 633546217 | PFL3 R4 | 942172835 | 3 |
| EPG L1 | 1447576662 | PFL3 L1 | 1258073453 | 32 |
| EPG L1 | 572870540 | PFL3 L1 | 1258073453 | 16 |
| EPG R1 | 5813014873 | PFL3 L1 | 1258073453 | 3 |
| EPG L2 | 912545106 | PFL3 L2 | 757694775 | 30 |
| EPG L2 | 697001770 | PFL3 L2 | 757694775 | 31 |
| EPG L1 | 572870540 | PFL3 L2 | 757694775 | 4 |
| EPG L1 | 1447576662 | PFL3 L2 | 757694775 | 7 |
| EPG R1 | 5813077544 | PFL3 R1 | 1134253374 | 21 |
| EPG R1 | 5813014873 | PFL3 R1 | 1134253374 | 21 |
| EPG R5 | 694920753 | PFL3 R5 | 911134017 | 22 |
| EPG R5 | 725951521 | PFL3 R5 | 911134017 | 10 |
| EPG R5 | 5813027103 | PFL3 R5 | 911134017 | 18 |
| EPG R6 | 942491983 | PFL3 R6 | 941132430 | 47 |
| EPG R5 | 725951521 | PFL3 R6 | 941132430 | 3 |
| EPG R6 | 941132434 | PFL3 R6 | 941132430 | 22 |
| EPG R5 | 5813027103 | PFL3 R6 | 941132430 | 5 |
| EPG R6 | 910438331 | PFL3 R6 | 941132430 | 28 |
| EPG R1 | 5813077544 | PFL3 R1 | 787374226 | 34 |
| EPG R1 | 5813014873 | PFL3 R1 | 787374226 | 22 |
| EPG R3 | 478375456 | PFL3 R3 | 912488890 | 21 |
| EPG R3 | 5813080838 | PFL3 R3 | 912488890 | 39 |
| EPG R3 | 541118908 | PFL3 R3 | 912488890 | 24 |
| EPG R5 | 5813027103 | PFL3 R5 | 910447181 | 17 |
| EPG R5 | 725951521 | PFL3 R5 | 910447181 | 27 |
| EPG R5 | 694920753 | PFL3 R5 | 910447181 | 14 |
| EPG R2 | 632544268 | PFL3 R2 | 1097718659 | 22 |
| EPG R2 | 695629525 | PFL3 R2 | 1097718659 | 33 |
| EPG R1 | 5813014873 | PFL3 R2 | 1097718659 | 3 |
| EPG R1 | 5813077544 | PFL3 R2 | 1097718659 | 8 |
| EPG L4 | 5813022281 | PFL3 L4 | 789130596 | 26 |
| EPG L4 | 665314820 | PFL3 L4 | 789130596 | 28 |
| EPG L5 | 819828986 | PFL3 L4 | 789130596 | 2 |
| EPG L4 | 634962055 | PFL3 L4 | 789130596 | 25 |
| EPG R6 | 910438331 | PFL3 R7 | 1200032115 | 2 |
| EPG R6 | 941132434 | PFL3 R7 | 1200032115 | 2 |
| EPG R6 | 942491983 | PFL3 R7 | 1200032115 | 4 |
| EPG R7 | 1002852791 | PFL3 R7 | 1200032115 | 40 |
| EPG R7 | 1034219901 | PFL3 R7 | 1200032115 | 24 |

| PreN | PreN ID | PostN | PostN ID | #syn |
|---|---|---|---|---|
| EPG R7 | 1002507159 | PFL3 R7 | 1200032115 | 35 |
| EPG L4 | 634962055 | PFL3 L5 | 880875927 | 3 |
| EPG L5 | 819828986 | PFL3 L5 | 880875927 | 27 |
| EPG L4 | 665314820 | PFL3 L5 | 880875927 | 3 |
| EPG L5 | 789126240 | PFL3 L5 | 880875927 | 10 |
| EPG L5 | 696362840 | PFL3 L5 | 880875927 | 13 |
| EPG L4 | 5813022281 | PFL3 L5 | 880875927 | 2 |
| EPG L5 | 789126240 | PFL3 L3 | 850855138 | 2 |
| EPG L4 | 5813022281 | PFL3 L3 | 850855138 | 10 |
| EPG L3 | 449438847 | PFL3 L3 | 850855138 | 2 |
| EPG L3 | 387364605 | PFL3 L3 | 850855138 | 22 |
| EPG L4 | 634962055 | PFL3 L3 | 850855138 | 30 |
| EPG L3 | 758419409 | PFL3 L3 | 850855138 | 7 |
| EPG L4 | 665314820 | PFL3 L3 | 850855138 | 13 |
| EPG L5 | 819828986 | PFL3 L5 | 911569552 | 24 |
| EPG L5 | 789126240 | PFL3 L5 | 911569552 | 4 |
| EPG L5 | 696362840 | PFL3 L5 | 911569552 | 11 |
| EPG L7 | 1035045015 | PFL3 L5 | 911569552 | 2 |
| EPG L6 | 912601268 | PFL3 L6 | 941939879 | 20 |
| EPG L5 | 789126240 | PFL3 L6 | 941939879 | 2 |
| EPG L6 | 788794171 | PFL3 L6 | 941939879 | 7 |
| EPG L5 | 696362840 | PFL3 L6 | 941939879 | 3 |
| EPG L7 | 1035045015 | PFL3 L6 | 941939879 | 2 |
| EPG L6 | 541870397 | PFL3 L6 | 941939879 | 28 |
| EPG L3 | 758419409 | PFL3 L3 | 912925080 | 28 |
| EPG L3 | 449438847 | PFL3 L3 | 912925080 | 25 |
| EPG L4 | 634962055 | PFL3 L3 | 912925080 | 2 |
| EPG L3 | 387364605 | PFL3 L3 | 912925080 | 31 |
| EPG R1 | 5813077544 | PFL3 R2 | 666994301 | 3 |
| EPG R2 | 632544268 | PFL3 R2 | 666994301 | 17 |
| EPG R1 | 5813014873 | PFL3 R2 | 666994301 | 2 |
| EPG R2 | 695629525 | PFL3 R2 | 666994301 | 21 |
| EPG L7 | 5813012006 | PFL3 L7 | 1004700437 | 15 |
| EPG L7 | 1035045015 | PFL3 L7 | 1004700437 | 18 |
| EPG L7 | 1004017998 | PFL3 L7 | 1004700437 | 7 |
| EPG L1 | 572870540 | PFL3 L1 | 944262351 | 16 |
| EPG L1 | 1447576662 | PFL3 L1 | 944262351 | 17 |
| EPG R1 | 5813077544 | PFL3 R1 | 666308023 | 24 |
| EPG L1 | 572870540 | PFL3 R1 | 666308023 | 6 |
| EPG L1 | 1447576662 | PFL3 R1 | 666308023 | 4 |
| EPG R1 | 5813014873 | PFL3 R1 | 666308023 | 17 |
| EPG L2 | 697001770 | PFL3 L2 | 1258686925 | 24 |
| EPG L1 | 572870540 | PFL3 L2 | 1258686925 | 7 |
| EPG L2 | 912545106 | PFL3 L2 | 1258686925 | 39 |
| EPG L1 | 1447576662 | PFL3 L2 | 1258686925 | 5 |
| EPG L1 | 1447576662 | PFL3 L1 | 851493896 | 29 |
| EPG R1 | 5813077544 | PFL3 L1 | 851493896 | 2 |
| EPG L1 | 572870540 | PFL3 L1 | 851493896 | 10 |

Table 1: **List of the synaptic connection considered to define the EPG-PFL3 connectome.**
PreN - Pre-synaptic neuron; PostN - Post-synaptic neuron; syn - Number of synapses

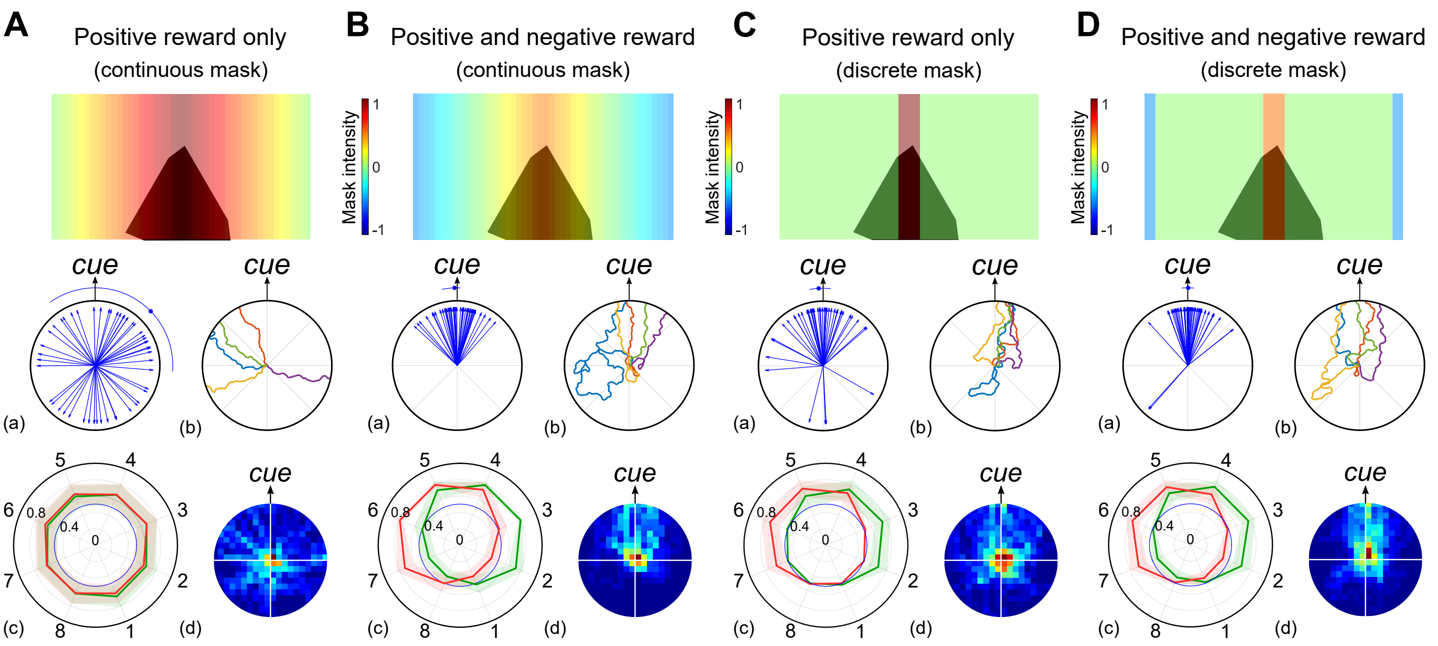


Figure S.2: **Innate attraction under the control of a visual reward signal.**
Simulations of the FB steering model (figure 6) using a reward signal provided by the visual processes to modify the EPG-PFL synapse weights. We created the visual input signal to the CX using different masks. Results for each panel include (a) the final path directions (n = 50 simulations), (b) examples of 5 simulation paths, (c) the averaged EPG-PFLs synapse weights and (d) a heat map of the agent's location.
**A.** Visual input to the FBs is equal to the sum of the visual units signal through a continuous proportional mask from 0 (rear units) to 1 (frontal units).
**B.** Visual input to the FBs is equal to the sum of the visual units signal through a continuous proportional mask from -0.5 (rear units) to 0.5 (frontal units).
**C.** Visual input to the FBs is equal to the sum of the visual units signal through a discrete mask equal to 0 (outside the 30° frontal area) or 1 (inside the 30° frontal area).
**D.** Visual input to the FBs is equal to the sum of the visual units signal through a discrete mask equal to 0 (outside the 30° frontal area and the 30° rear area), 0.5 (inside the 30° rear area) or 1 (inside the 30° frontal area).

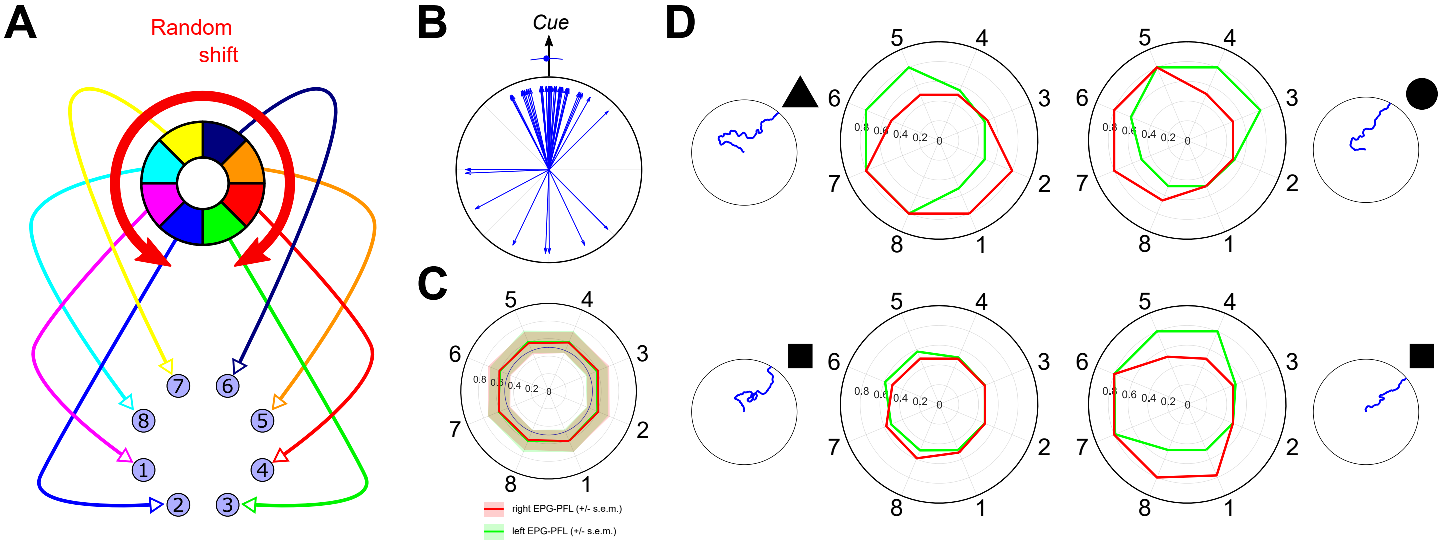


Figure S.3: **Using an offset compass does not compromise the model function.**
**A.** The connectivity between the visual units and the E-PGs has been shifted by a random value. The bump produced therefore expresses a constant offset different for every simulation.
**B.** The innate attraction behaviour is not affected by the offset applied on the EB bump.
**C.** Overall EPG-PFL synapse weights. The lines indicate the mean value for each EPG-PFL couple (red for the right side and green for the left) and the shaded area the standard mean deviation (s.e.m.).
**D.** 4 individual experiment paths and the associated EPG-PFL synapse weights generated during the simulation (red for the right side and green for the left).

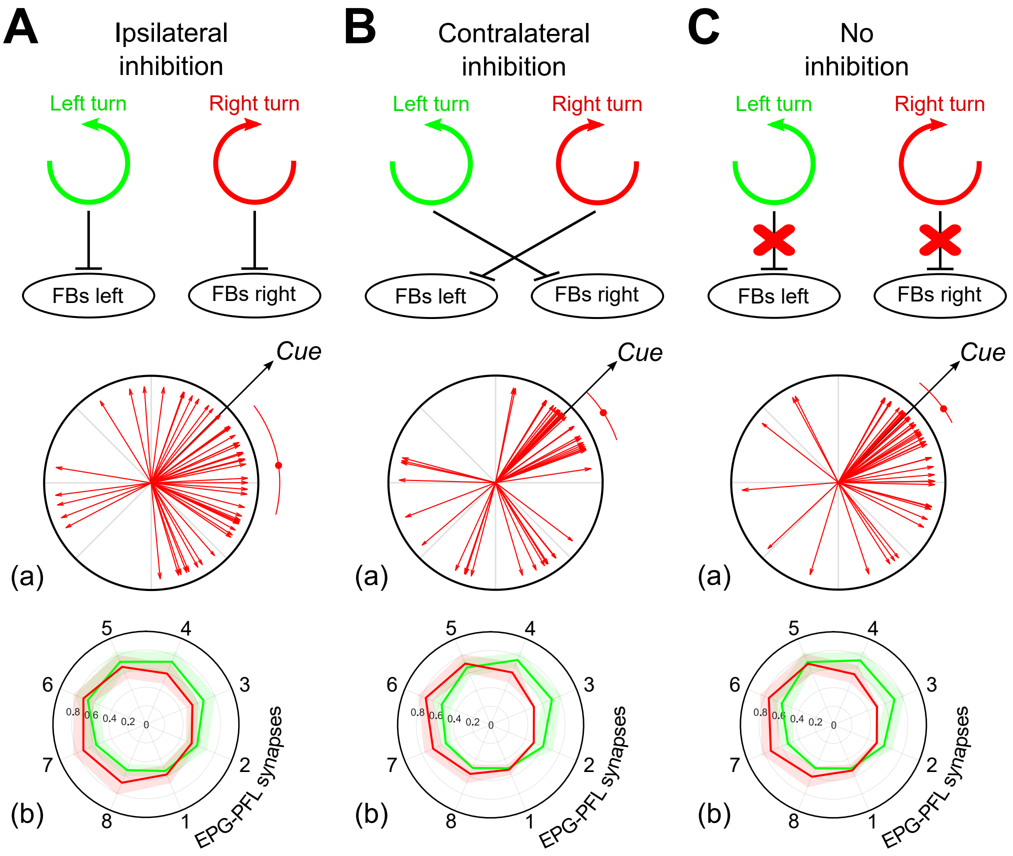


Figure S.4: **Impact of the self motion integration in the model.**
**A-B-C.** Model simulations with a combination of innate attraction and visual memory (MBs) as presented in figure 10A & B. (a) Final direction vectors for 50 simulations. The arc represent the median (dot) ± 95% C.I. obtained via bootstrap (rep = 10000). (b) Averaged right (red) and left (green) EPG-PFL synapse weights (shaded area: ±*s.d.*) obtain during 50 simulations.
**A.** Ipsilateral inhibitory circuit from the self-motion signal to the FBns. This corresponds to the circuit presented in figure 6.
**B.** Contralateral inhibitory circuit from the self-motion signal to the FBns.
**C.** Circuit without any integration of the self-motion by the FBns layer.

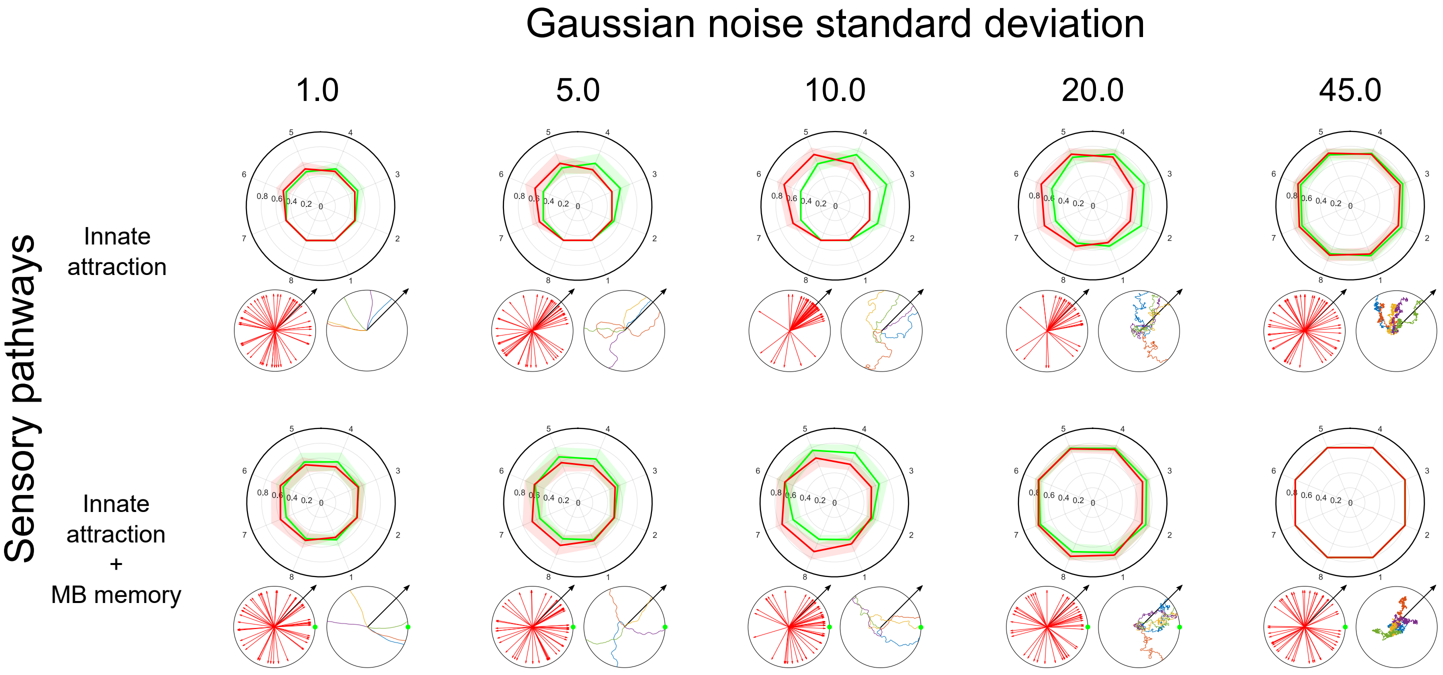


Figure S.5: **Impact of the steering noise on the model performance.**
The influence of the noise standard deviation applied to the steering is tested in the case of the innate attraction alone ($\omega_{MB} = 0$) and combined with the visual memory ($\omega_{Vin}$ and $\omega_{MB}$ set as in figure 10B).

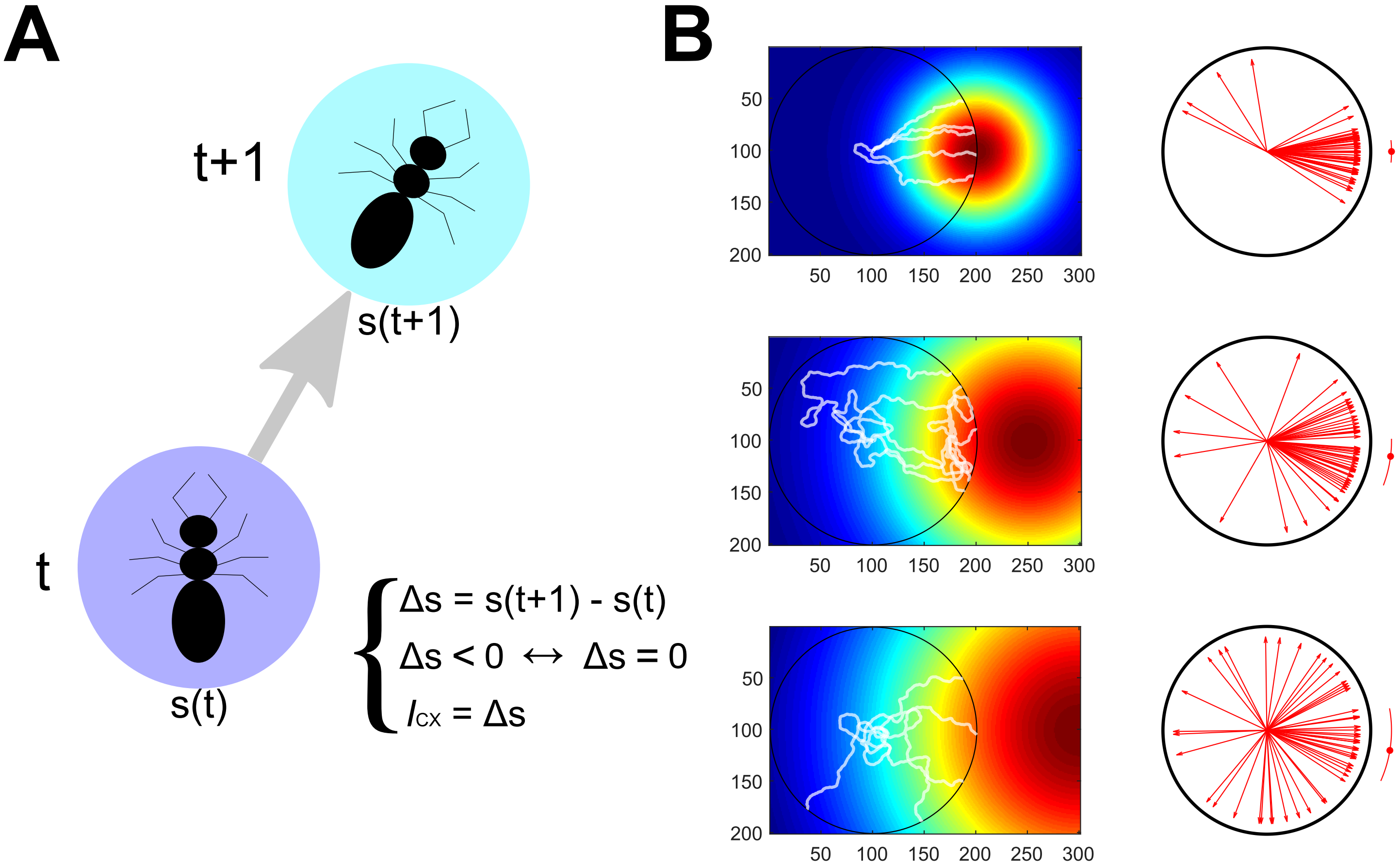


Figure S.6: **Gradient ascent properties, potential for the olfactory pathway.**
**A.** The reward signal ($I_{CX}$, input to the CX) is defined here as the difference ($\Delta s$) of the concentration in sensory input $[s(t)]$. Negative input, decrease of the concentration are not considered here ($\Delta s < 0 \leftrightarrow \Delta s = 0$). The value of $\Delta$ is multiplied by a free parameter $\omega_{Olf}$ to adapt roughly the amplitude to the same range as the visual pathways (innate and learned). During the simulations the visual compass is maintain, and keep it's function to modulate the EPG-PFL3 synaptic weights, thanks to a landmark randomly positioned around the arena.
**B.** Simulations with 3 different sources positioned at 100, 150 and 200 lu. from the arena centre creating a gaussian gradient (respectively s.d. = 50, 100 and 150 lu.). Left panels show the gradient shape and 5 paths example (white lines). Right panels show the final direction of the simulations (n = 50) and the median (dot) ± 95% C.I. (arc) obtained by bootstrap (rep = 10000).

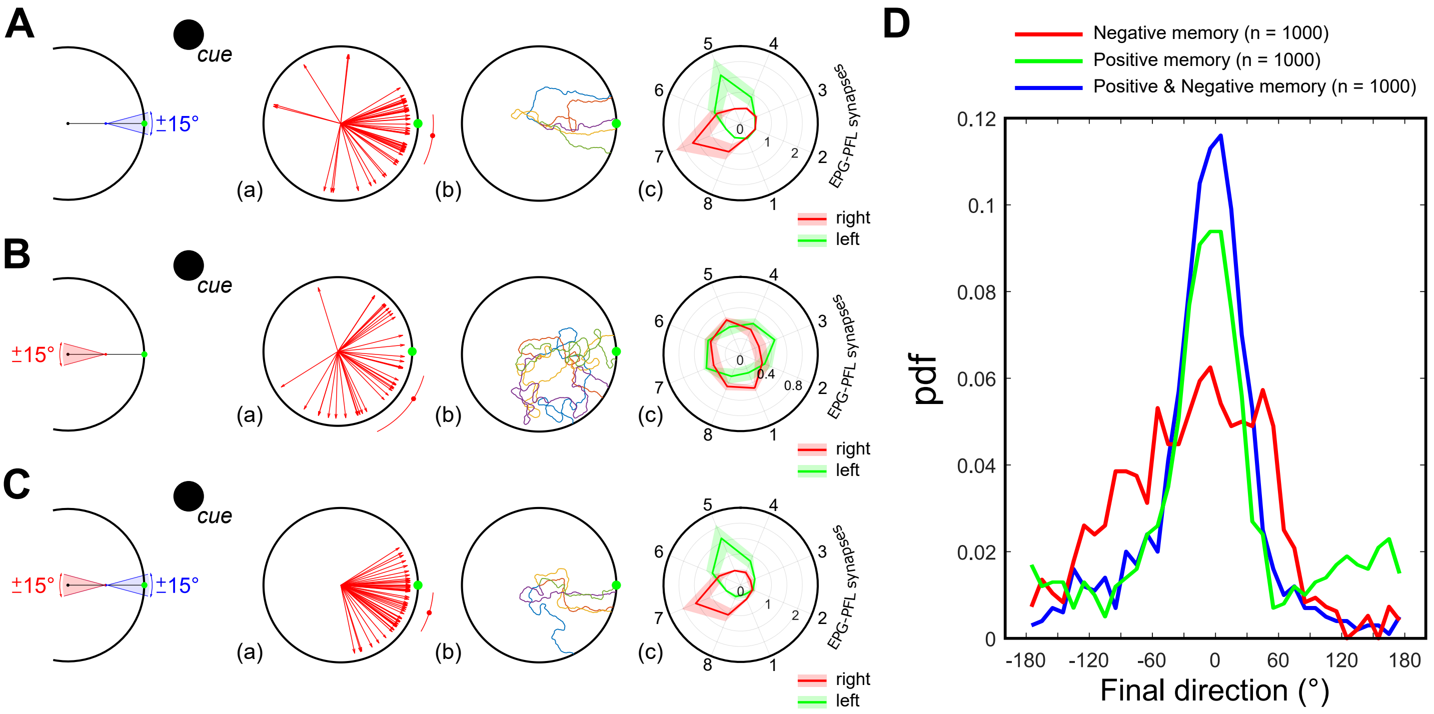


Figure S.7: **Influence of a attractive-repulsive MB views memory on the model.**
**A-B-C.** (a) Final direction vectors of 50 simulations. The red arc indicates the median (red dot) 95% C.I. obtained by bootstrap ($n_{rep} = 10000$). (b) 5 path examples. (c) Averaged right (red) and left (green) EPG-PFLs synapse weights (shaded area: $\pm s.d.$). For these simulations, in contrary to the other results presented in the paper, synapses weights were not capped to 0.8.
**A.** Simulations with attractive views memory only acquired facing (±15°, blue span) the feeder (green dot). This correspond to the same simulations as presented in figure 9.
**B.** Simulations with repulsive views memory only. MBON value is multiplied by -1 before connecting the FB neurons. Repulsive views are acquired in the same manner than the attractive one while facing 180° ± 15° (red span) away from the feeder (green dot).
**C.** Simulations with both attractive and repulsive views memory. Each memory use the very same subset of KC neurons but is processed on a different MBON (positive for attraction, negative for repulsion).
**D.** Probability density function of the final directions for 1000 simulations in the 3 conditions (Attractive memory, positive memory and attractive & positive memory).
